# The Functional Nanopore Screen: A Versatile High-throughput Assay to Study and Engineer Protein Nanopores in *Escherichia coli*

**DOI:** 10.1101/2021.04.20.440580

**Authors:** Wadim Weber, Markus Roeder, Tobias Probanowski, Jie Yang, Helal Abujubara, Heinz Koeppl, Alesia Tietze, Viktor Stein

## Abstract

Nanopores comprise a versatile class of membrane proteins that carry out a range of key physiological functions and are increasingly developed for different biotechnological applications. Yet, a capacity to study and engineer protein nanopores by combinatorial means has so far been hampered by a lack of suitable assays that combine sufficient experimental resolution with throughput. Addressing this technological gap, the Functional Nanopore (FuN) screen now provides a quantitative and dynamic read-out of nanopore assembly and function in *E. coli*. The assay is based on genetically-encoded fluorescent protein sensors that resolve the nanopore-dependent influx of Ca^2+^ across the inner membrane of *E. coli*. Illustrating its versatile capacity, the FuN screen is first applied to dissect the molecular features that underlie the assembly and stability of nanopores formed by the S^21^68 holin. In a subsequent step, nanopores are engineered by recombining the transmembrane module of S^21^68 with different ring-shaped oligomeric protein structures that feature defined hexa-, hepta- and octameric geometries. Library screening highlights substantial plasticity in the ability of the S^21^68 transmembrane module to oligomerize in alternative geometries while the functional properties of the resultant nanopores can be fine-tuned through the identity of the connecting linkers. Overall, the FuN screen is anticipated to facilitate both fundamental studies and complex nanopore engineering endeavors with many potential applications in biomedicine, biotechnology and synthetic biology.

## Introduction

Nanopores comprise a versatile class of membrane proteins that form aqueous channels across cellular membranes to facilitate the passage of polar and charged molecules across an otherwise impermeable barrier. As part of their natural function, nanopores underlie a range of key physiological processes such as cell lysis^1^ or the permeation of antibiotics^2^. Recent years also witnessed the development of nanopores for different biotechnological applications^3^ most prominently in the context of biosensing^4^, DNA sequencing^4^ and single molecule studies^5^. To this end, a range of different protein nanopores scaffolds have been assessed including oligomeric toxins such as αHLA, MspA, FraC and ClyA^6–13^, bacterial outer membrane proteins such as FhuA and OmpG^14–18^ and self-assembling membrane peptides^19–22^. In addition, protein nanopores could recently be engineered by computational means either using a consensus design approach^22^ or *de novo* based on first principles^23–25^.

Despite an increasing number of nanopores being studied and subject to sophisticated engineering endeavors, their experimental characterization remains technically challenging. In fact, progress has been hampered for a number of reasons: Firstly, given their capacity to form aqueous channels across cellular membranes, nanopores are toxic and therefore cannot be easily expressed in recombinant hosts. Secondly, nanopores frequently comprise large, integral membrane proteins that rely on lipid environments for their functional expression and characterization. Thirdly, protein nanopores may also depend on accessory factors to efficiently insert and assemble in cellular membranes.

The expression of nanopores is thus frequently limited to reconstituted lipid bilayers *in vitro*^26^ while functional studies – for example by electrophysiological^27^ or optical means^28^ – are typically restricted in terms of throughput. In a few limited instances, high-throughput screening and selection systems such as liposome display^8^ or hemolytic assays^12^ have been devised, but are currently limited by technically challenging protocols. For instance, liposome display relies on artificial, yet fragile cell-like compartments generated by *in vitro* compartmentalization that do not fully recapitulate critical features such as the membrane potential^8^. Alternatively, hemolytic assays are restricted to specific types of nanopores, in particular, bacterial toxins that critically depend on the membrane composition of red blood cells to assemble.

Addressing these limitations, the functional nanopore (FuN) screen now provides a scalable, quantitative and time-resolved read-out of nanopore function in *E. coli*. Following its experimental validation, the FuN screen is used to dissect the molecular features that underlie the assembly and functional integrity of nanopores formed by the prototypical S^21^68 holin^29,30^. In further efforts, a C-terminally truncated variant of the S^21^68 holin is recombined with ring-shaped hexa-, hepta- and octameric protein structures probing the plasticity of the S^21^68 transmembrane module to oligomerize while fine-tuning the permeability of the resultant nanopore assemblies solely *via* the connecting linkers. Overall, the FuN screen should thus greatly facilitate both fundamental studies and complex nanopore engineering endeavors.

## Results

### FuN Screen Experimental Implementation

To assay protein nanopores in live cells, a reporter-based genetic screening system was developed (**Figure 1**). The assay relies on genetically-encoded Ca^2+^ indicators for optical imaging (GE-COs)^31^ to resolve the nanopore-dependent influx of Ca^2+^ across the inner membrane and thus provide a time-resolved optical read-out of nanopore function in *E. coli*. To ensure independent control over the expression of the reporter and the nanopore, transcriptional units were placed on separate plasmids while their transcription was initiated with *E. coli* and T7 RNA polymerases under the control of propionate and IPTG-inducible promoters, respectively.

**Figure 1.**
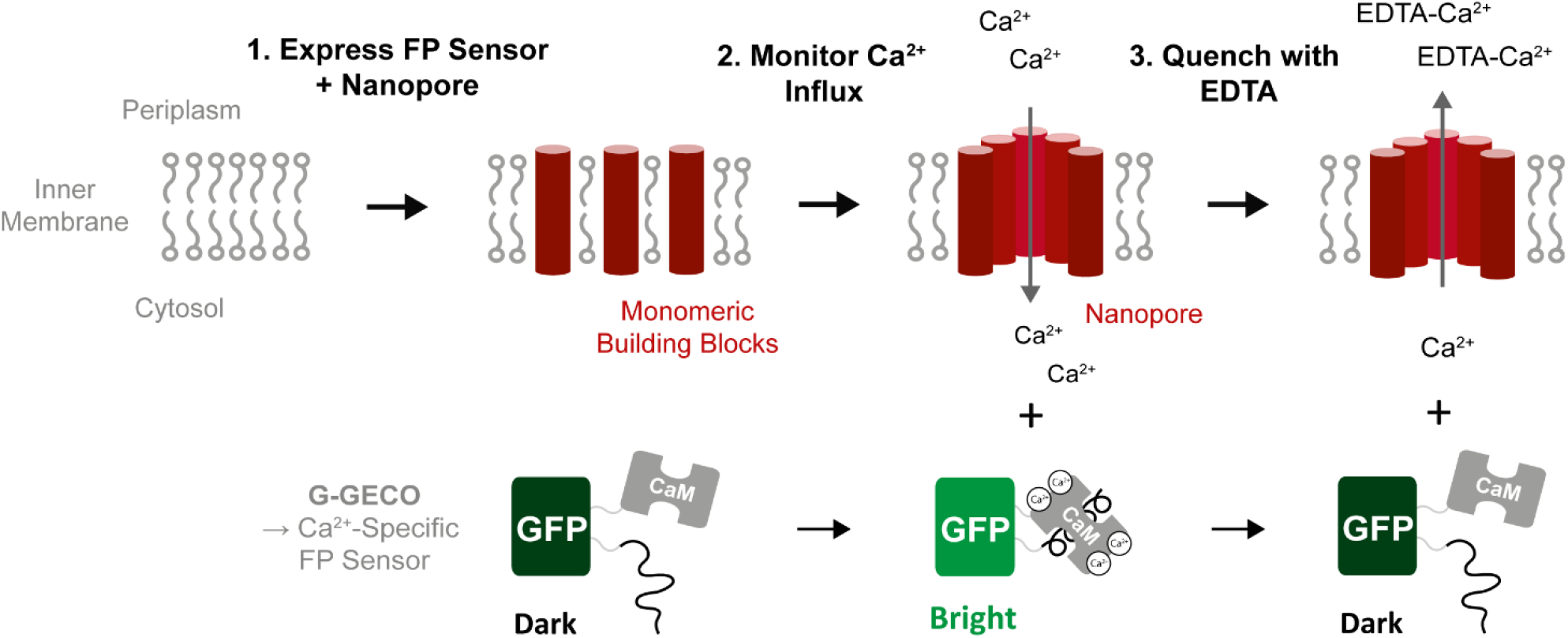
Schematic of the FuN screen assay. Following functional expression of a nanopore, Ca^2+^ ions flow across the inner membrane into the cytoplasm of *E. coli*. The influx of Ca^2+^ ions is experimentally resolved using a genetically-encoded Ca^2+^-specific fluorescent protein (FP) sensor that provides a quantitative and time-resolved optical read-out of nanopore function in *E. coli*. In a subsequent step, the fluorescent signal is quenched with EDTA to provide complementary information on the structural and function integrity of a protein nanopore independent of its insertion, activation and assembly kinetics.

With a set of expression constructs established, it was first examined whether the fluorescent signal associated with a Ca^2+^-dependent influx is specific to the functional properties of different nanopores and ion channels. To this end, S^21^68 holin^29,30^, the T4 holin^32^, the K_CV_NTS^33^ and the transmembrane module of the H^+^-permeable BM2 channel^34^ were employed as controls. Both, the T4 holin and the S^21^68 holin derive from bacteriophages where they initiate cell lysis by forming nm- to µm-sized pores in the inner membrane of *E. coli*^35^. In contrast, K_CV_NTS and BM2 derive from Chlorella and Influenza B virus and are selective for K^+^ and H^+^ ions providing suitable negative controls. To afford a quantitative measure, the resultant fluorescent signal was empirically fit to Equation S1 where the time T_½_ required to reach half the maximum signal provides a measure for nanopore formation in *E. coli* (**Figure 2A)**. In addition, the OD_600_ was monitored independently to assess the functional expression of individual nanopores and its impact on cellular integrity. Furthermore, to balance toxicity associated with leaky expression in the absence of IPTG while achieving a strong quantitative fluorescent signal, it was critical to adjust the expression of individual nanopores and ion channels through tailored T7 promoters^36,37^.

**Figure 2.**
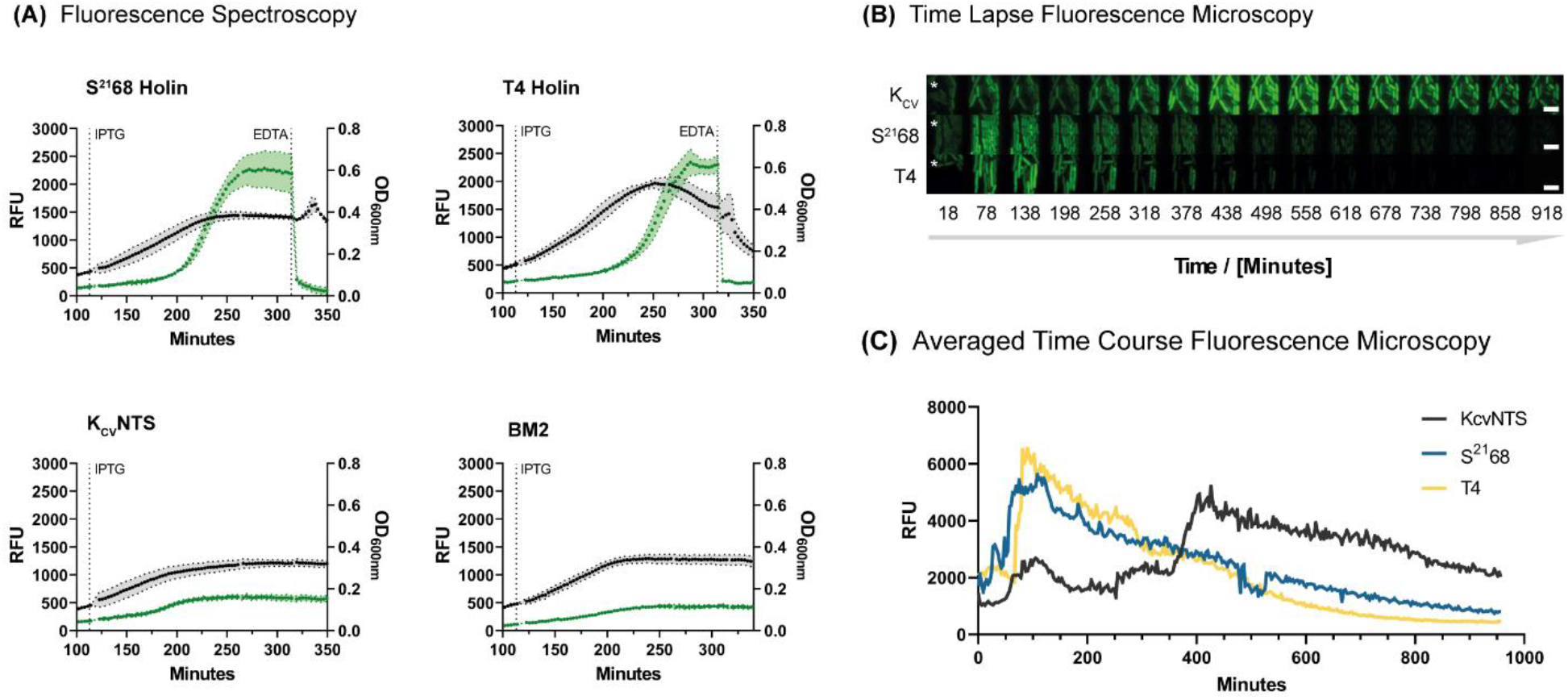
Experimental validation of the FuN screen. **(A)** The expression of S^21^68, T4 holin, K_CV_NTS and BM2 is induced with IPTG and the developing fluorescent signal monitored in a microplate reader. The expression of S^21^68 and the T4 holin, but not K_CV_NTS and BM2 triggers a strong increase in the fluorescent signal demonstrating that an influx of Ca^2+^ correlates with the functional properties of a nanopore and ion channel. The fluorescent signal is quenched by adding EDTA; **(B)** Correlating the developing fluorescent signal for S^21^68 holin, the T4 holin and K_CV_NTS channel with cellular physiology at single cell resolution following growth in a microfluidic incubation chamber and imaging with time lapse fluorescence microscopy. Cellular integrity remains intact upon expression of S^21^68 and the K_CV_NTS channel, but not the T4 holin; **(C)** Quantitative comparison of the developing fluorescent signal averaged across the microfluidic incubation chamber. In contrast to K_CV_NTS, the expression of the S^21^68 and the T4 holin trigger a rapid increase in the fluorescent signal demonstrating the specificity of the FuN screen. Nanopores and ion channels were expressed from the pCtrl2.

Crucially, the expression of S^21^68 and the T4 holin triggered a strong increase in the fluorescent signal while the K^+^-selective K_CV_NTS channel and the H^+^-permeable BM2 channel did not (**Figure 2A**). Subsequent addition of EDTA triggered a rapid drop in the fluorescent signal for both the S^21^68 and the T4 holin (**Figure 2A**). Differences were however observed in terms of the OD_600_ which dropped sharply for the T4 holin but declined more gradually for S^21^68. These differences can be attributed to different effects on cellular integrity as the T4 holin forms large µm-sized holes that cause a cell to disintegrate^38^ while the S^21^68 holin forms smaller, defined nm-sized nanopores leaving the cell intact for a prolonged period of time^39^. Supporting this notion, the expression of the T4 holin releases a significant amount of fluorescence in the supernatant indicating cell rupture (**Figure S1**). In contrast, the fluorescent signal remains in the pellet following expression of the S^21^68 holin or K_CV_NTS indicating cellular integrity remains intact (**Figure S1**).

To examine how the functional expression of a nanopore affects the integrity of the cell and how it correlates with the developing fluorescent signal, the expression of the S^21^68, the T4 holin and the K_CV_NTS channel was further examined by time-resolved fluorescence microscopy at single cell resolution in a microfluidic incubation chamber (**Figure 2B**). In all cases, cells rapidly stopped to divide while the fluorescent signal averaged across the microfluidic incubation chamber (**Figure 2C**) matched the course of the fluorescent signal observed by fluorescence spectroscopy in microtitre plates (**Figure 2A**). Notably, the expression of the T4 holin and S^21^68 triggered a strong early increase in the fluorescent signal within 80-100 min before gradually fading (as a result of bleaching). In comparison, only a very late increase in the fluorescent signal was observed for K_CV_NTS at 380 min. This could either be due to a breakdown of membrane homeostasis or a low, non-specific influx of Ca^2+^ into the cell, which eventually triggers a Ca^2+^-dependent fluorescent signal in the cell. Furthermore, in terms of cellular integrity, cellular structures disappear for the T4 holin as it forms µm-sized holes that trigger cell lysis and enable the G-GECO to diffuse out of the cell. In contrast, cellular structures remain intact for prolonged periods of time following expression of S^21^68 and K_CV_NTS.

### Dissecting S^21^68 Nanopore Activation

With elementary protocols established, the FuN screen was applied to dissect the molecular features that underlie the formation of nanopores by the S^21^68 holin (**Figure 3**). The current mechanism hypothesizes that S^21^68 along with its cognate anti-holin S^21^71, which features an additional three amino acid cytoplasmic anchor at its N-terminus, initially accumulate as inactive, anti-parallel α-helical homodimers in the inner membrane of *E. coli*^40^. Upon reaching a critical concentration, the N-terminal transmembrane domain, termed TMD1, flips across the inner membrane and assumed to initiate the assembly of heptameric nanopores^29^ (**Figure 3B**). Yet, little is known which molecular features control the translocation of TMD1 across the inner membrane and its contribution to the structural and functional integrity of nanopores formed by the S^21^68 holin.

**Figure 3.**
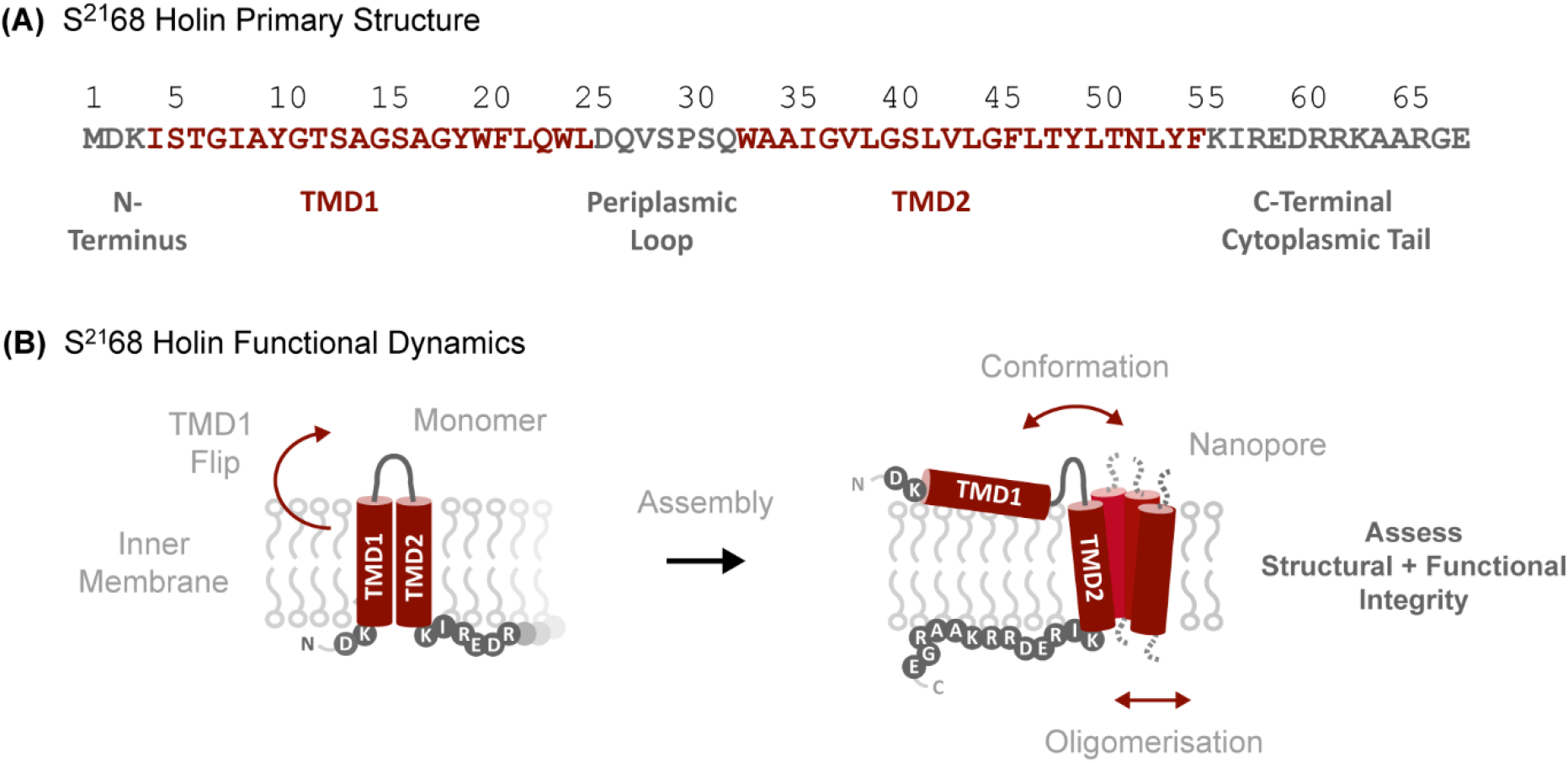
Functional assembly of the S^21^68 holin. **(A)** Primary structure of the S^21^68 holin. Transmembrane regions are denoted in red while cytoplasmic and periplasmic regions are denoted in grey. **(B)** Simplified schematic overview of the dynamic features that underlie the formation of nanopores by the S^21^68 holin. Upon reaching a critical concentration in the inner membrane, the N-terminal transmembrane domain (TMD1) flips across the inner membrane to initiate nanopore assembly.

Thus, to gain a more quantitative understanding of the molecular features underlying this dynamic assembly process, the two most N-terminal residues D2 and K3 were substituted with 17 different amino acids and their effect on the formation of S^21^68 nanopores quantified in terms of the time to reach half the maximum signal T_½_ (**Figure 4A** and **Figure S2**). To prevent artefacts arising through alternative start codons and disulfide bridges, Met and Cys were omitted from the library. Substitutions generally had a greater effect on K3 relative to D2. Notably, all substitutions in K3 except for Arg significantly accelerate the formation of S^21^68 nanopores suggesting that positively charged side chains anchor the N-terminus *via* electrostatic interactions with the negatively charged phospholipid head groups in the cytoplasm and thus delay translocation of TMD1 across the inner membrane (**Figure 4A**). Conversely, hydrophobic and negatively charged side chains assisted by the negative membrane potential facilitate translocation of TMD1 across the bilayer. In contrast, substitution of D2 generally exerted a lesser effect on the propensity of S^21^68 to form nanopores (**Figure S2**) – i.e. the hydrophobic substitutions Ile, Leu and Phe slightly accelerated nanopore formation as the lipophilicity and thus ability of the S^21^68 N-terminus to pass the hydrophobic core of the lipid bilayer is enhanced. In contrast, the polar uncharged residues Ser, Thr, Asn and Gln slowed the formation of S^21^68 nanopores effectively removing the charge associated with D2 that otherwise drives the translocation of TMD1 across the membrane. Combined, these trends readily conform with the ‘positive-inside’ and ‘negative-outside’ rules that have been empirically determined to rationalize the folding and insertion of transmembrane proteins^41–43^ while highlighting strong positional effects of individual residues regulating the ability of TMD1 to translocate across the inner membrane and initiate the assembly of nanopores.

**Figure 4.**
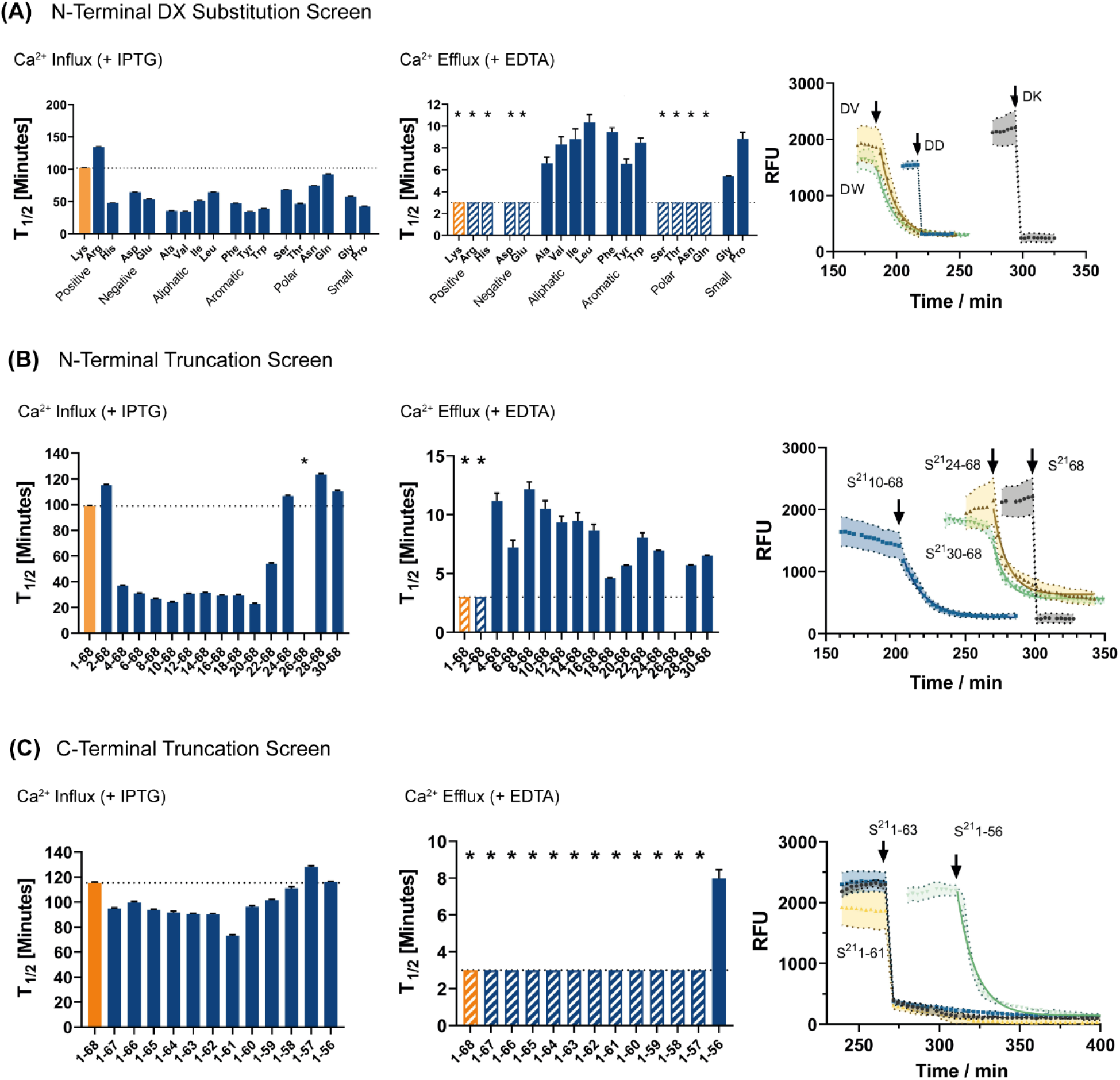
Dissecting the molecular features mediating the assembly and stability of S^21^68 nanopores. First, **(A)** K3 was substituted with 17 different amino acids, then, **(B)** TMD1 truncated in increments of two amino acids, and finally, **(C)** the C-terminus truncated in increments of one amino acid. The propensity to form stable protein nanopores is assessed in terms of the time T_½_ to reach half the maximum signal following addition of IPTG and the half-life T_½_ of the fluorescent signal following addition of EDTA. A set of representative kinetic traces is shown for EDTA quench experiments highlighting a highly variable signal decay for select mutants. For mutants denoted with *, the half-life T_½_ is less than 3 min and thus below the temporal resolution of the assay. Black arrows denote the time point of adding 5 mM EDTA. S^21^68 variants were expressed from pCtrl2.

Further, the functional properties of S^21^68 nanopores were analyzed following the addition of EDTA when the fluorescent signal saturated and cells stopped to divide (**Figure 4A**). Considering nanopores already formed but are no longer expressed at this stage, the signal becomes independent of the expression, insertion and activation kinetics and primarily depends on the structural and functional integrity of nanopores in the assembled state. Again, striking differences were observed for substitutions in K3 (**Figure 4A**) but not D2 (**Figure S2**). This time the ability to quench the fluorescent signal with EDTA strongly correlated with the polarity of side chains (**Figure 4A**). Notably, hydrophilic residues behave comparable to K3 as the fluorescent signal was rapidly quenched upon addition of EDTA. In contrast, hydrophobic residues showed a marked deceleration in Ca^2+^ efflux. In particular, a decreased Ca^2+^ efflux scaled with the hydrophobicity of aliphatic side chains (Ala > Val > Ile > Leu) suggesting that hydrophobic substitutions in K3 draw TMD1 deeper into the membrane and thus adversely impact the structural and functional integrity of S^21^68 nanopores in the assembled state. Overall, systematic N-terminal substitutions thus highlight a critical role for K3 in timing the assembly, but also in maintaining the structural and functional integrity of S^21^68.

### Delineating Minimal S^21^68 Pore Forming Motifs

In a subsequent set of experiments, S^21^68 was systematically truncated in increments of two amino acids to identify the molecular features that are necessary and sufficient to form functional nanopores. The propensity of truncated variants to form nanopores was then quantified in terms of the time T_½_ required to reach half the maximum signal (**Figure 4B** and **Figure S3**). Notably, the ability to form nanopores decreased slightly upon removal of D2 (see S^21^2-68), but was quickly enhanced upon deletion of K3 and I4 (see S^21^4-68) further underpinning the inhibitory role of K3 which is partially offset by the negative charge of D2.

A very strong propensity to form nanopores was then maintained until deletion of the aromatic patch YWFLQW (see S^21^24-68) and partially regained upon removal of the hydrophilic loop LDQVSPSQ connecting TMD1 with TMD2 (see S^21^30-68). Considering a decreased fluorescent signal could either arise from a decelerated ability to form, or stem from a compromised structural and functional integrity in the assembled state, truncation mutants were subject to EDTA quench experiments independent of their expression, insertion and activation kinetics (**Figure 4B**).

Notably, once K3 was deleted, the ability of truncation mutants to form functional nanopores was reduced as judged by decreased Ca^2+^ efflux rates indicating an increase in conformational heterogeneity, changes in the oligomerization state or a combination thereof. Conversely, an enhanced capacity of truncation mutants to form nanopores as judged by the time to reach half the maximum signal upon addition of IPTG is primarily enabled by an enhanced capacity of the N-terminus to translocate across the inner membrane, especially once K2 is deleted. This becomes particularly obvious when comparing the course of the fluorescent signal of S^21^68 and S^21^2-68 with S^21^24-68 and S^21^30-68 which display comparable kinetics of formation, but different stabilities in the assembled state as judged by EDTA quench rates (**Figure 4B**). Furthermore, truncations variants with N-terminal aromatic residues, especially S^21^18-68, partially recover in stability suggesting a contribution of aromatic stacking interactions between TMD1 and the lipid bilayer in stabilizing S^21^68 nanopores in the assembled state.

In contrast to the N-terminus, truncation of the C-terminus does not negatively impact the formation of S^21^68 nanopores and turns out largely dispensable for assembly (**Figure 4C**). Only truncation to K56 causes an increase in Ca^2+^ efflux rates highlighting a critical need to appropriately anchor the C-terminus of S^21^68 at the cytoplasmic interface of the membrane through electrostatic interactions with the phospholipid headgroups. This was further corroborated in substitution experiments (**Figure S4**). Here, only Arg could functionally replace K56, even enhancing Ca^2+^ efflux indicating a partial recovery in the structural and functional integrity of the S^21^56 variant while the titratable His and negatively charged Glu conferred significantly reduced and hydrophobic amino acids only very limited functionality. Combined these trends also quantitatively recapitulate the distribution of amino acids in transmembrane domains^43^ highlighting the experimental resolution of the FuN screen.

### S^21^68 TMD1 is Essential to Form Uniformly Stable Nanopores

To gain a more quantitative understanding of N-terminal truncations, the S^21^68 holin along with the S^21^10-68, S^21^24-68, and S^21^30-68 variants were chemically synthesized and electrophysiologically characterized in reconstituted vertical lipid bilayer membranes (**Figure 5**). Electrophysiological measurements demonstrate that full-length S^21^68 holin forms nanopores of unitary conductance providing further evidence that it assembles into defined nm-sized nanopores following insertion and activation in the membrane (**Figure 5A**). Furthermore, distinct conductive states were observed for S^21^10-68 and S^21^24-68 (**Figure 5B, 5C**) while the shortest truncation S^21^30-68 only yielded poorly defined pores with characteristic spikes reflecting the transient nature of this minimal variant (**Figure 5D**). Electrophysiological measurements thus confirm that TMD2 of S^21^68 is necessary and sufficient to form conductive nanopores – albeit only very transient ones. This observation further emphasizes the stabilizing contribution of the very N-terminus as judged by the length and uniformity of the conductive states of the full length S^21^68 holin compared to the shorter S^21^10-68, S^21^24-68 and especially the S^21^30-68 truncations. Furthermore, deletion of the aromatic, hydrophobic patch YWFLQW, which is entirely absent in S^21^30-68, results in highly transient nanopores indicating stabilizing contacts between the aromatic residues in TMD1 and the membrane.

**Figure 5.**
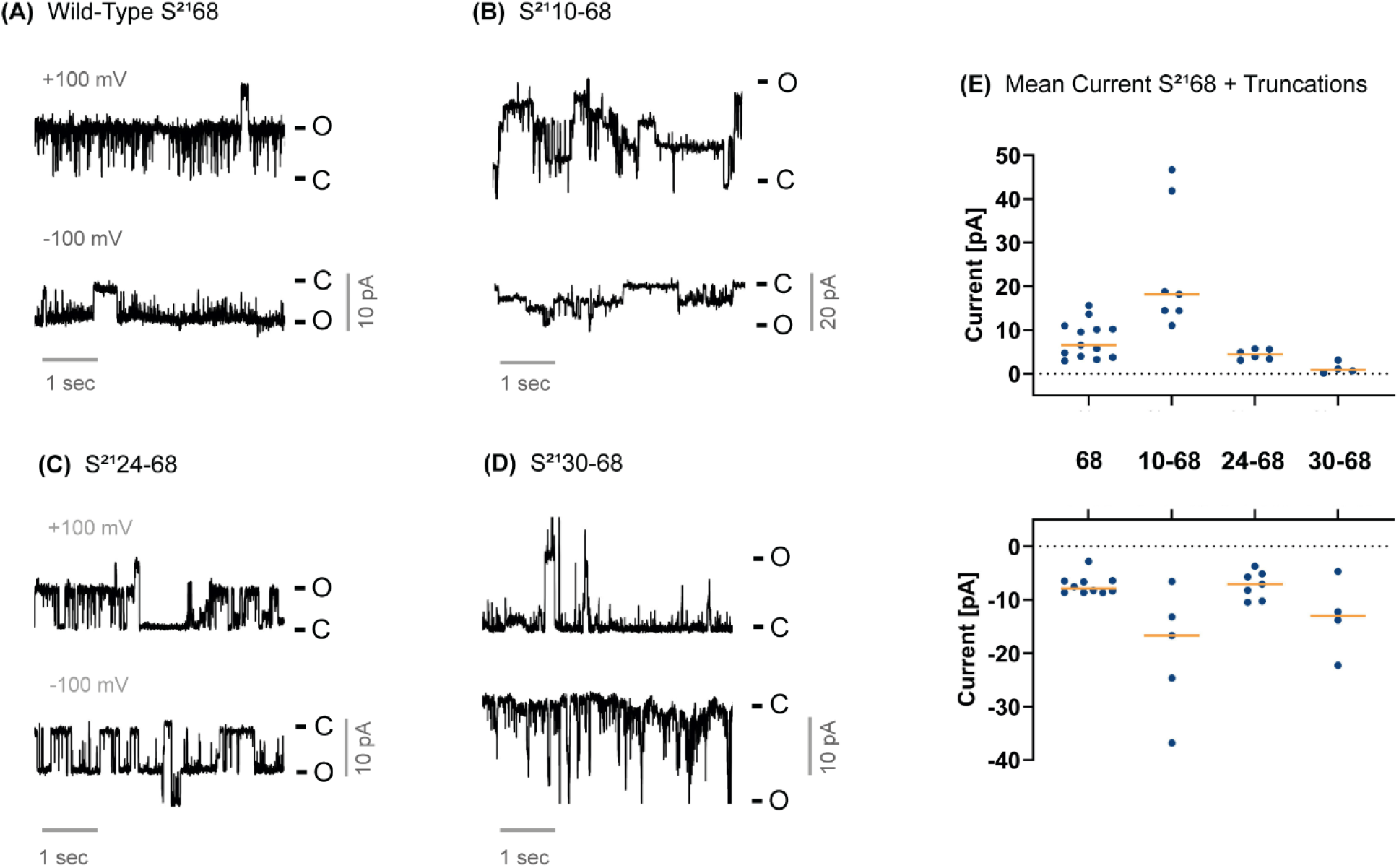
Electrophysiological characterization of the S^21^68 holin and a select number of truncation mutants. **(A-D)** Summary of representative electrical recordings of different S^21^68 variants; The least and most conductive states are denoted with C and O; **(E)** The average current of individual truncation mutants correlates with the time T_½_ to reach half the maximum signal observed in the FuN screen as S^21^10-68 enables on average the greatest conductivity. The average current of a single 5 sec -100 mV or +100 mV protocol is indicated by a blue dot and orange lines mark the median.

Furthermore, electrical recordings demonstrate a good correlation with the fluorescent signal observed in the FuN screen highlighting its physiological relevance. Notably, S^21^10-68 also displayed the strongest propensity to assemble into functional nanopores in reconstituted lipid bilayer membranes as judged by the average conductivity (**Figure 5E**). In significant parts, this is driven by the capacity of truncation variants to efficiently insert into membranes which is also reflected by an accelerated time to reach half the maximum signal following addition of IPTG (**Figure 4B**). Conversely, only the full length S^21^68 holin is capable of forming uniformly stable nanopores for prolonged periods of time (**Figure 5A**) underpinning rapid EDTA quench rates (**Figure 4B**). Combined, these results demonstrate the exquisite quantitative and temporal resolution that can be achieved with the FuN screen in dissecting the dynamic features that underlie the assembly of protein nanopores directly in the context of genetically tractable *E. coli*.

### Engineering Recombinant Nanopore Assemblies

Finally, a set of recombinant nanopores was engineered by recombining the truncated S^21^56 variant with ring-shaped oligomeric protein structures and linkers of varying length and flexibility^44,45^ (**Figure 6A**). Ring-shaped oligomeric protein structures were drawn from members of the Lsm protein family AfSm2^46^, AfSm1^47^ and ScLsm3^48^ featuring defined hexa-, hepta- and octameric geometries. Visibly, oligomerization accelerated the assembly of nanopores as judged by the time to reach half the maximum signal following addition of IPTG, and, on average, turned out most compatible with the native heptameric conformation of the S^21^68 holin (**Figure 6C**). Surprisingly, recombination with hexameric and octameric protein structures also yielded highly functional nanopores emphasizing substantial functional plasticity in TMD2 albeit with a greater dependence on the identity of the connecting linkers as judged by the average spread across linker libraries that was most pronounced for octameric ScLsm3 (**Figure 6D**) and to a lesser extent affected heptameric AfSm2 (**Figure 6B**).

**Figure 6.**
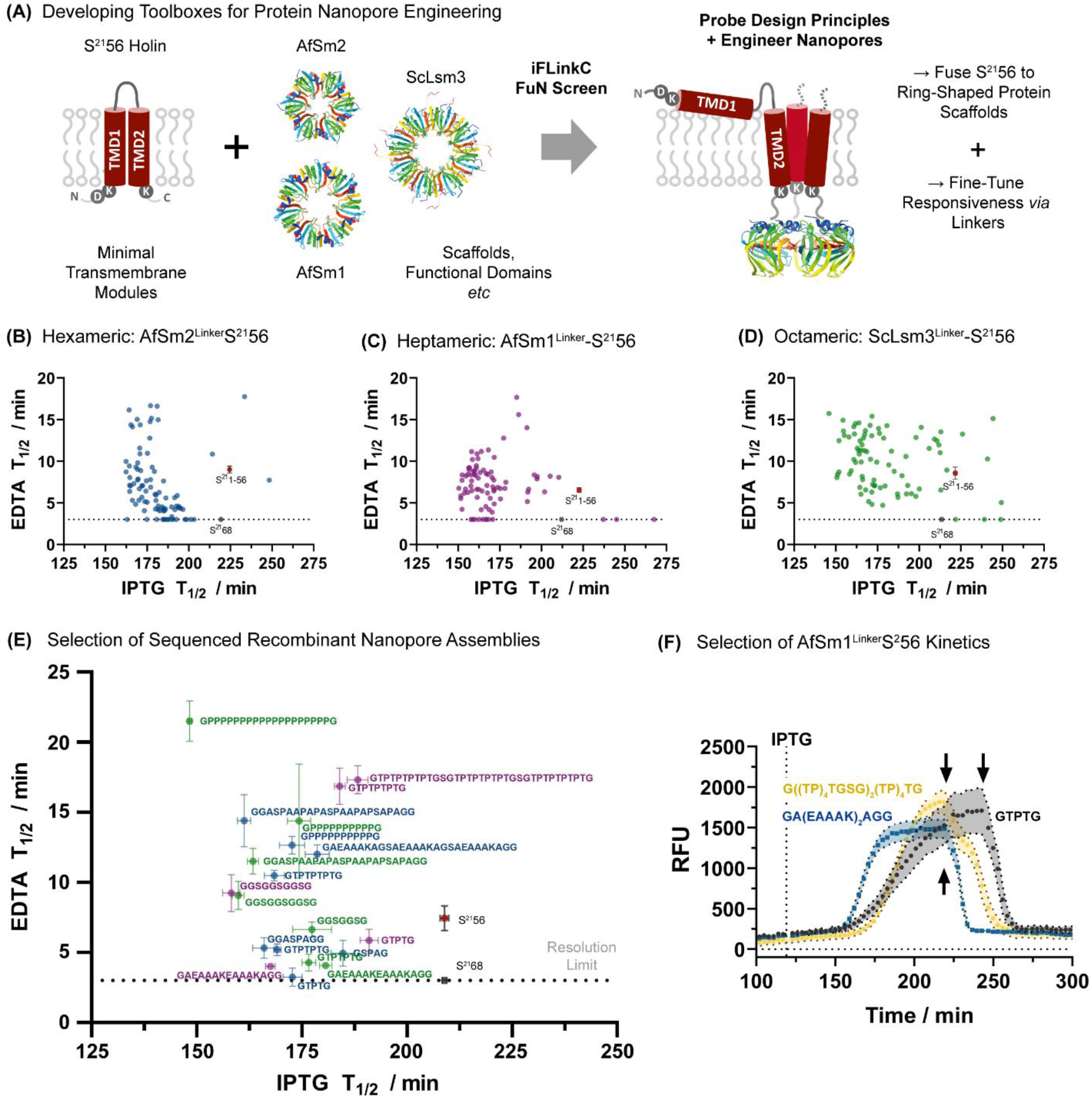
Engineering recombinant protein nanopore assemblies. **(A)** Recombinant nanopores are engineered through systematic recombination of a pore-forming S^21^56 transmembrane module to ring-shaped oligomeric protein structures. Ring-shaped oligomeric protein structures feature defined hexa-, hepta- and octameric geometries while fusion is achieved through combinatorial linker libraries; **(B-D)** Summary of AfSm1, AfSm2 and ScLsm3 library screens. The functional properties of recombinant nanopores are assessed in terms of the time to reach half the maximal signal upon addition of IPTG and the half-life of the fluorescent signal upon addition of EDTA; **(E)** Summary of assembly (+ IPTG) and efflux rates (+ EDTA) for a select number of sequenced linker variants in the context of different hexa-, hepta- and octameric geometries highlighted in blue, purple and green (n = 4); Oligomerization strongly accelerates nanopore assembly while linkers modulate Ca^2+^ efflux; Reference measurements of the truncated S^21^56 variant and wild-type S^21^68 are included for every microtiter plate measurement; **(F)** Kinetic traces of a select number of S^21^56-AfSm1 linker variants. Recombinant nanopores assemblies and thereof composed linker libraries were expressed from the pCtrl3. PDB Codes: AfSm2 (1LJO), AfSm1 (3BW1) and ScLsm3 (1I4K).

Conversely, the ability of recombinant nanopores to mediate an efflux of Ca^2+^ independent of their expression, insertion and assembly kinetics critically depends on the identity of the connecting linkers yielding nanopores with enhanced, neutral and worse Ca^2+^ efflux rates relative to the truncated S^21^56 variant (**Figure 6B-D**). Re-assaying a selection of sequence-verified nanopore variants across the three libraries revealed a preference for short semi-flexible linkers of 8 amino acids or less and a limited number of structured linkers (**Figure 6E, 6F**). Notably, in the context of heptameric AfSm1, an α-helical linker performed best enhancing assembly and accelerating Ca^2+^ efflux while flexible TP- and GS-rich linkers of >9 were associated with low Ca^2+^ efflux. Similarly, in the context of hexameric AfSm2, the most efficient Ca^2+^ efflux was associated with short semi-flexible linker motifs comprising 8 residues or less while linkers of 9 residues or more including highly structured once were generally associated with decreased Ca^2+^ efflux rates (**Figure 6E**). Comparable trends were observed for octameric ScLsm3. Recombination of ring-shaped oligomeric protein structures with a pore-forming transmembrane module thus readily yields functional nanopores while their permeability can be fine-tuned *via* the connecting linkers.

## Discussion

Addressing a critical need in nanopore engineering, nanopores can now be studied and engineered by means of the FuN screen. The assay relies on the nanopore-dependent influx of Ca^2+^ into *E. coli* which is resolved with genetically-encoded Ca^2+^-specific FP sensors. Crucially, the assay recapitulates the functional properties of different nanopores and ion channels as it is specific for Ca^2+^ while K^+^ and H^+^-permeable channels such as K_CV_NTS and BM2 do not trigger a signal. Furthermore, the FuN screen carries a number of unique advantages. Firstly, its read-out is compatible with different optical assay formats, for instance, in microtitre plates for medium throughput, colony on-plate for high-throughput, or fluorescence microscopy at single cell resolution in a microfluidic incubation chamber. Secondly, all components are genetically encoded and therefore do not require external reagents. Thirdly, experimental protocols are simple and can be readily realized in any molecular biology lab. Fourthly, the FuN screen is quantitative and capable of resolving the dynamic features that underlie the assembly and function of nanopores and potentially other membrane proteins mediating the passage of ions and solutes across the inner membrane. Fifthly, given the availability of FP sensors, the FuN screen is readily scalable to study and engineer the permeation of distinct analytes across the inner membrane of *E. coli* according to the functional properties of different nanopores and ion channels.

In a proof-of-concept, the FuN screen is used to dissect the molecular features underlying the formation of S^21^68 nanopores. In this regard, early genetic and recent biophysical studies suggest TMD2 is necessary and sufficient to form functional nanopores while the N-terminus times its activation and assembly^49–53^. Systematic truncation of TMD1 complemented by electrophysiological measurements with chemically synthesized peptides support this notion – yet, highlight that TMD2 is only capable of forming transient nanopores. This becomes particularly evident for S^21^30-68 which is entirely devoid of TMD1 while portions of TMD1 successively contribute to its structural and functional integrity. Notably, inclusion of an aromatic patch, which forms part of TMD1, partially stabilizes S^21^68 nanopores while uniformly conductive nanopores are only observed for the full-length S^21^68 holin.

Further insight concerns the activation of S^21^68 which highlights carefully crafted N-terminal features and position-specific effects: in particular, K3 turns out critical for timing as it delays activation as both deletion and substitution of K3 (with the exception of Arg) rapidly accelerate nanopore formation no longer anchoring the N-terminus of TMD1 in the cytosol *via* electrostatic interactions with the phospholipid headgroups. Yet, surprisingly, K3 also contributes to the structural and functional integrity of S^21^68 nanopores which is negatively impacted by hydrophobic, but not negatively charged or polar substitutions. This implies a generic need for anchoring the N-terminus of S^21^68 in the periplasm in order to generate homogenously stable nanopores.

Finally, the FuN screen is applied in the construction of recombinant nanopore assemblies. To this end, a truncated S^21^56 variant is recombined with ring-shaped oligomeric protein structures and linkers of variable length and flexibility. In the context of the S^21^68 holin, recombination yields functional evidence for a preferred heptameric geometry of TMD2^30^ – yet, also demonstrates substantial plasticity in its ability to oligomerize in non-native hexa- and octameric geometries. In addition, the functional properties of recombinant nanopores can be fine-tuned *via* the identity of the connecting linkers highlighting an opportunity for linker engineering in nanopore engineering modulating their assembly efficiency or fine-tuning their permeability. Equally, the combinatorial assembly of recombinant nanopores expands the repertoire of ring-shaped oligomeric protein structures that are available for nanopore engineering^54^.

## Conclusions

The FuN screen now provides a potent assay to study and engineer nanopores in *E. coli* testing and informing on the design principles of natural as well as artificially engineered nanopores. In particular, it combines throughput with quantitative resolution in the context of a genetically-tractable organism. The FuN screen is thus anticipated to provide a powerful complementary methodology to high-resolution biophysical techniques based on electrophysiological^27^ and optical read-outs^28^ that are limited to measurements in reconstituted artificial lipid bilayers.

Applications are diverse and include the development of therapeutic agents based on pore-forming membrane peptides, equipping nanopores with bespoke sensory and regulatory features or engineering their ability to insert into lipid bilayers independent of cellular chaperones, for instance, in the context of artificial cells. Conversely, systematic recombination of transmembrane modules with ring-shaped protein structures highlights the potential of the FuN screen in probing the molecular features that underlie the formation of oligomeric protein complexes. As far as the S^21^68 holin is concerned, its systematic genetic dissection highlights complex sequence-structure-function relationships underlying its functional assembly that motivate further high-resolution structural and computational studies.

## Author Contributions

**WW** designed experiments, performed experiments, analyzed data and wrote the manuscript. **MR** performed experiments. **TP** performed experiments and analyzed data. **JY** chemically synthesized S^21^68 holin variants. **HA** chemically synthesized S^21^68 holin variants. **HK** assisted with microfluidic experimental set up and analysis. **AT** directed the chemical synthesis of S^21^68 holin variants. **VS** conceived and directed study, analyzed data and wrote the manuscript. All authors have given approval to the final version of the manuscript.

## Notes

VS, WW and MR are co-inventors on a patent application submitted by the TU Darmstadt that covers the described methodology.

## Acknowledgements

The authors acknowledge support by Christine Pleitner, Theresa Wörmann and Sarah Schmitt assisting with experimental efforts, Gerhard Thiel for helpful comments and suggestions concerning the electrophysiological characterization of the S^21^68 holin and thereof derived variants, and Yolanda Schaerli for consulting on microfluidic chip designs.

## References

(1) Peraro, M. D.,, Van Der Goot, F. G. Pore-Forming Toxins: Ancient, but Never Really out of Fashion. Nature Reviews Microbiology. 2016.

(2) Vergalli, J.; Bodrenko, I. V.; Masi, M.; Moynié, L.; Acosta-Gutiérrez, S.; Naismith, J. H.; Davin-Regli, A.; Ceccarelli, M.; van den Berg, B.; Winterhalter, M.; et al. Porins and Small-Molecule Translocation across the Outer Membrane of Gram-Negative Bacteria. Nature Reviews Microbiology. 2020.

(3) Ayub, M.; Bayley, H. Engineered Transmembrane Pores. Curr. Opin. Chem. Biol. 2016, 34, 117–126.

(4) Wang, S.; Zhao, Z.; Haque, F.; Guo, P. Engineering of Protein Nanopores for Sequencing, Chemical or Protein Sensing and Disease Diagnosis. Current Opinion in Biotechnology. 2018.

(5) Movileanu, L. Interrogating Single Proteins through Nanopores: Challenges and Opportunities. Trends in Biotechnology. 2009.

(6) Gu, L. Q.; Braha, O.; Conlan, S.; Cheley, S.; Bayley, H. Stochastic Sensing of Organic Analytes by a Pore-Forming Protein Containing a Molecular Adapter. Nature 1999.

(7) Braha, O.; Walker, B.; Cheley, S.; Kasianowicz, J. J.; Song, L.; Gouaux, J. E.; Bayley, H. Designed Protein Pores as Components for Biosensors. Chem. Biol. 1997.

(8) Fujii, S.; Matsuura, T.; Sunami, T.; Kazuta, Y.; Yomo, T. In Vitro Evolution of α-Hemolysin Using a Liposome Display. Proc. Natl. Acad. Sci. U. S. A. 2013, 110 (42), 16796–16801.

(9) Derrington, I. M.; Butler, T. Z.; Collins, M. D.; Manrao, E.; Pavlenok, M.; Niederweis, M.; Gundlach, J. H. Nanopore DNA Sequencing with MspA. Proc. Natl. Acad. Sci. U. S. A. 2010.

(10) Wloka, C.; Mutter, N. L.; Soskine, M.; Maglia, G. Alpha-Helical Fragaceatoxin C Nanopore Engineered for Double-Stranded and Single-Stranded Nucleic Acid Analysis. Angew. Chemie - Int. Ed. 2016, 55 (40), 12494–12498.

(11) Huang, G.; Willems, K.; Soskine, M.; Wloka, C.; Maglia, G. Electro-Osmotic Capture and Ionic Discrimination of Peptide and Protein Biomarkers with FraC Nanopores. Nat. Commun. 2017.

(12) Soskine, M.; Biesemans, A.; De Maeyer, M.; Maglia, G. Tuning the Size and Properties of ClyA Nanopores Assisted by Directed Evolution. J. Am. Chem. Soc. 2013, 135 (36), 13456–13463.

(13) Stoddart, D.; Ayub, M.; Höfler, L.; Raychaudhuri, P.; Klingelhoefer, J. W.; Maglia, G.; Heron, A.; Bayley, H. Functional Truncated Membrane Pores. Proc. Natl. Acad. Sci. U. S. A. 2014, 111 (7).

(14) Thakur, A. K.; Movileanu, L. Real-Time Measurement of Protein–Protein Interactions at Single-Molecule Resolution Using a Biological Nanopore. Nat. Biotechnol. 2019.

(15) Thakur, A. K.; Movileanu, L. Single-Molecule Protein Detection in a Biofluid Using a Quantitative Nanopore Sensor. ACS Sensors 2019.

(16) Liu, Z.; Ghai, I.; Winterhalter, M.; Schwaneberg, U. Engineering Enhanced Pore Sizes Using FhuA Δ1-160 from E. Coli Outer Membrane as Template. ACS Sensors 2017.

(17) Chen, M.; Khalid, S.; Sansom, M. S. P.; Bayley, H. Outer Membrane Protein G: Engineering a Quiet Pore for Biosensing. Proc. Natl. Acad. Sci. U. S. A. 2008.

(18) Sanganna Gari, R. R.; Seelheim, P.; Liang, B.; Tamm, L. K. Quiet Outer Membrane Protein G (OmpG) Nanopore for Biosensing. ACS Sensors 2019.

(19) Lella, M.; Kamilla, S.; Jain, V.; Mahalakshmi, R. Molecular Mechanism of Holin Transmembrane Domain i in Pore Formation and Bacterial Cell Death. ACS Chem. Biol. 2016, 11 (4), 910–920.

(20) Krishnan, S.; Satheesan, R.; Puthumadathil, N.; Kumar, K. S.; Jayasree, P.; Mahendran, K. R. Autonomously Assembled Synthetic Transmembrane Peptide Pore. J. Am. Chem. Soc. 2019.

(21) Lella, M.; Mahalakshmi, R. Engineering a Transmembrane Nanopore Ion Channel from a Membrane Breaker Peptide. J. Phys. Chem. Lett. 2016.

(22) Mahendran, K. R.; Niitsu, A.; Kong, L.; Thomson, A. R.; Sessions, R. B.; Woolfson, D. N.; Bayley, H. A Monodisperse Transmembrane α-Helical Peptide Barrel. Nat. Chem. 2017.

(23) Vorobieva, A. A.; White, P.; Liang, B.; Horne, J. E.; Bera, A. K.; Chow, C. M.; Gerben, S.; Marx, S.; Kang, A.; Stiving, A. Q.; et al. De Novo Design of Transmembrane β Barrels. Science (80-.). 2021, 371 (6531).

(24) Xu, C.; Lu, P.; Gamal El-Din, T. M.; Pei, X. Y.; Johnson, M. C.; Uyeda, A.; Bick, M. J.; Xu, Q.; Jiang, D.; Bai, H.; et al. Computational Design of Transmembrane Pores. Nature 2020, 585 (7823).

(25) Scott, A. J.; Niitsu, A.; Kratochvil, H. T.; Lang, E. J. M.; Sengel, J. T.; Dawson, W. M.; Mahendran, K. R.; Mravic, M.; Thomson, A. R.; Brady, R. L.; et al. Constructing Ion Channels from Water-Soluble α-Helical Barrels. Nat. Chem. 2021.

(26) Shen, H. H.; Lithgow, T.; Martin, L. L. Reconstitution of Membrane Proteins into Model Membranes: Seeking Better Ways to Retain Protein Activities. International Journal of Molecular Sciences. 2013.

(27) Gutsmann, T.; Heimburg, T.; Keyser, U.; Mahendran, K. R.; Winterhalter, M. Protein Reconstitution into Freestanding Planar Lipid Membranes for Electrophysiological Characterization. Nat. Protoc. 2015, 10 (1).

(28) Heron, A. J.; Thompson, J. R.; Cronin, B.; Bayley, H.; Wallace, M. I. Simultaneous Measurement of Ionic Current and Fluorescence from Single Protein Pores. J. Am. Chem. Soc. 2009, 131 (5).

(29) Park, T.; Struck, D. K.; Deaton, J. F.; Young, R. Topological Dynamics of Holins in Programmed Bacterial Lysis. Proc. Natl. Acad. Sci. U. S. A. 2006.

(30) Pang, T.; Savva, C. G.; Fleming, K. G.; Struck, D. K.; Young, R. Structure of the Lethal Phage Pinhole. Proc. Natl. Acad. Sci. U. S. A. 2009, 106 (45).

(31) Kalko, E. K. V; Dukas, R.; Ratcliffe, J. M.; Teeling, E. C.; Haven, N.; Fattu, J. M.; Bates, M. E.; Simmons, J. a; Riquimaroux, H.; Surlykke, A.; et al. An Expanded Palette of Genetically Encoded Ca2+ Indicators. Science (80-.). 2011, 333, 1888–1891.

(32) Ramanculov, E.; Young, R. Genetic Analysis of the T4 Holin: Timing and Topology. Gene 2001.

(33) Pagliuca, C.; Goetze, T. A.; Wagner, R.; Thiel, G.; Moroni, A.; Parcej, D. Molecular Properties of Kcv, a Virus Encoded K+ Channel. Biochemistry 2007, 46 (4).

(34) Mandala, V. S.; Loftis, A. R.; Shcherbakov, A. A.; Pentelute, B. L.; Hong, M. Atomic Structures of Closed and Open Influenza B M2 Proton Channel Reveal the Conduction Mechanism. Nat. Struct. Mol. Biol. 2020, 27 (2).

(35) Reddy, B. L.; Saier, M. H. Topological and Phylogenetic Analyses of Bacterial Holin Families and Superfamilies. Biochim. Biophys. Acta - Biomembr. 2013.

(36) Imburgio, D.; Rong, M.; Ma, K.; McAllister, W. T. Studies of Promoter Recognition and Start Site Selection by T7 RNA Polymerase Using a Comprehensive Collection of Promoter Variants. Biochemistry 2000, 39 (34).

(37) Dubendorff, J. W.; Studier, F. W. Creation of a T7 Autogene. Cloning and Expression of the Gene for Bacteriophage T7 RNA Polymerase under Control of Its Cognate Promoter. J. Mol. Biol. 1991, 219 (1).

(38) Tran, T. A. T.; Struck, D. K.; Young, R. Periplasmic Domains Define Holin-Antiholin Interactions in T4 Lysis Inhibition. J. Bacteriol. 2005, 187 (19).

(39) Park, T.; Struck, D. K.; Dankenbring, C. A.; Young, R. The Pinholin of Lambdoid Phage 21: Control of Lysis by Membrane Depolarization. J. Bacteriol. 2007, 189 (24).

(40) Young, R. Bacteriophage Holins: Deadly Diversity. In Journal of Molecular Microbiology and Biotechnology; 2002; Vol. 4.

(41) Baker, J. A.; Wong, W. C.; Eisenhaber, B.; Warwicker, J.; Eisenhaber, F. Charged Residues next to Transmembrane Regions Revisited: “Positive-inside Rule” Is Complemented by the “Negative inside Depletion/Outside Enrichment Rule.” BMC Biol. 2017, 15 (1).

(42) von Heijne, G. The Distribution of Positively Charged Residues in Bacterial Inner Membrane Proteins Correlates with the Trans-Membrane Topology. EMBO J. 1986, 5 (11).

(43) Elazar, A.; Weinstein, J.; Biran, I.; Fridman, Y.; Bibi, E.; Fleishman, S. J. Mutational Scanning Reveals the Determinants of Protein Insertion and Association Energetics in the Plasma Membrane. Elife 2016, 5 (JANUARY2016).

(44) Gräwe, A.; Ranglack, J.; Weyrich, A.; Stein, V. IFLinkC : An Iterative Functional Linker Cloning Strategy for the Combinatorial Assembly and Recombination of Linker Peptides with Functional Domains. Nucleic Acids Res. 2020, 1–11.

(45) Gräwe, A.; Ranglack, J.; Weyrich, A.; Stein, V. Synthetic Protein Switches: Combinatorial Linker Engineering with IFLinkC. In Methods in Enzymology; 2021; Vol. 647.

(46) Törö, I.; Basquin, J.; Teo-Dreher, H.; Suck, D. Archaeal Sm Proteins Form Heptameric and Hexameric Complexes: Crystal Structures of the Sm1 and Sm2 Proteins from the Hyperthermophile Archaeoglobus Fulgidus. J. Mol. Biol. 2002, 320 (1).

(47) Törö, I.; Thore, S.; Mayer, C.; Basquin, J.; Séraphin, B.; Suck, D. RNA Binding in an Sm Core Domain: X-Ray Structure and Functional Analysis of an Archaeal Sm Protein Complex. EMBO J. 2001, 20 (9).

(48) Naidoo, N.; Harrop, S. J.; Sobti, M.; Haynes, P. A.; Szymczyna, B. R.; Williamson, J. R.; Curmi, P. M. G.; Mabbutt, B. C. Crystal Structure of Lsm3 Octamer from Saccharomyces Cerevisiae: Implications for Lsm Ring Organisation and Recruitment. J. Mol. Biol. 2008, 377 (5).

(49) Steger, L. M. E.; Kohlmeyer, A.; Wadhwani, P.; Bürck, J.; Strandberg, E.; Reichert, J.; Grage, S. L.; Afonin, S.; Kempfer, M.; Görner, A. C.; et al. Structural and Functional Characterization of the Pore-Forming Domain of Pinholin S2168. Proc. Natl. Acad. Sci. U. S. A. 2020, 117 (47).

(50) Ahammad, T.; Drew, D. L.; Sahu, I. D.; Serafin, R. A.; Clowes, K. R.; Lorigan, G. A. Continuous Wave Electron Paramagnetic Resonance Spectroscopy Reveals the Structural Topology and Dynamic Properties of Active Pinholin S2168 in a Lipid Bilayer. J. Phys. Chem. B 2019, 123 (38).

(51) Ahammad, T.; Drew, D. L.; Khan, R. H.; Sahu, I. D.; Faul, E.; Li, T.; Lorigan, G. A. Structural Dynamics and Topology of the Inactive Form of S21Holin in a Lipid Bilayer Using Continuous-Wave Electron Paramagnetic Resonance Spectroscopy. J. Phys. Chem. B 2020, 124 (26).

(52) Ahammad, T.; Drew, D. L.; Sahu, I. D.; Khan, R. H.; Butcher, B. J.; Serafin, R. A.; Galende, P.; McCarrick, R. M.; Lorigan, G. A. Conformational Differences Are Observed for the Active and Inactive Forms of Pinholin S21 Using Deer Spectroscopy. J. Phys. Chem. B 2020, 124 (50).

(53) Pang, T.; Fleming, T. C.; Pogliano, K.; Young, R. Visualization of Pinholin Lesions in Vivo. Proc. Natl. Acad. Sci. U. S. A. 2013, 110 (22).

(54) Ho, C. W.; Meervelt, V. Van Tsai, K. C.; De Temmerman, P. J.; Mast, J.; Maglia, G. Engineering a Nanopore with Co-Chaperonin Function. Sci. Adv. 2015, 1 (11).

