## Supporting Information for "The Functional Nanopore Screen: A Versatile High-throughput Assay to Study and Engineer Protein Nanopores in *Escherichia coli*"

**This PDF file contains:**

Supporting Methods
Supporting Results
Figures S1-S4
SI References

### Supporting Methods

*Materials*

The DNA coding for the G-GECO fluorescent sensor, different nanopores including ion channels and thereof derived libraries was either purchased from public repositories (Addgene) or commercially synthesized as ssDNA (Sigma-Aldrich) or gBlocks (IDT DNA Technologies). Constructs were generally cloned in conventional DH10B (New England Biolabs) before being transformed, expressed and assayed in BL21 (DE3) (New England Biolabs). The expression of the reporter and the nanopore was mediated from two separate transcriptional units that were placed on separate plasmids with compatible origins of replication while their transcription was initiated with *E. coli* and T7 RNA polymerases under the control of propionate and IPTG-inducible promoters, respectively.

*Recombinant DNA Work*

Constructs to express different FP sensors and nanopores were generated by conventional restriction-digestion-ligation cloning, Gibson Assembly[1] or iFLinkC[2]. Expression constructs were generally based on the pPro24[3,4], pCtrl2 or pCtrl3. The latter two expression constructs are based on the pACYCT2[5] backbone that have been optimised for the functional expression protein nanopores in *E. coli*.

*The FuN Screen Assay*

Different protein nanopores and the G-GECO fluorescent sensor were expressed under the control of IPTG and propionate-inducible promoters from two separate expression plasmids based on the pCtrl2, pCtrl3 and pPro24 backbones, respectively. Except for BM2, protein nanopores were expressed from a weakened T7 promoter[6]. To reduce nanopore expression in the basal state, a LacI mutant W220F with enhanced repression properties was used[7]. Briefly, plasmids coding for the G-GECO[8] reporter and the nanopore of interest were co-transformed into BL21 (DE3) cells and plated on LB agar plates supplemented with 100 µg/mL ampicillin (AMP) and 25 µg/mL chloramphenicol (CHL). For spectroscopic screening, individual colonies were used to inoculate 300 µL lysogeny broth (LB) medium supplemented with 100 µg/mL AMP and 25 µg mg/mL CHL and grown overnight in 96 deep-well plates at 37 ˚C and 1300 rpm. The following day 3.5 µL cell suspension was added to 196.5 µL LB supplemented with 25 mM sodium propionate pH 8.0, 100 µg/mL AMP and 25 µg/mL CHL in 96-well black microtitre plates with transparent bottom. Plates were incubated in a spectroscopic microtiter plate reader (TECAN Spark) at 30 ˚C with fluorescence and $OD_{600}$ measurement every 3 minutes while being shaken at 180 rpm. Following 110 min of incubation, nanopore expression was initiated by adding 0.5 mM IPTG. The development of the fluorescent signal was measured through Ex. 480 ± 10 nm and Em. 525 ± 10 nm (Gain 60) along with the $OD_{600}$ to assess cellular integrity. To quantify assembly kinetics, the fluorescent signal was fit to **Eq. S1** through non-linear regression and the time to reach half the maximum signal $T_{½}$ used as a measure for the assembly of functional nanopores.

**Eq. S1** $$\boldsymbol{Y_{(x)} = Bottom + \frac{Top}{1+e^{\frac{-(x-T_{1/2})}{Slope}}}}$$

For EDTA quench experiments, 5 mM EDTA was added approximately 15 min after the fluorescent signal saturated and cells stopped to divide as judged by the $OD_{600}$. The decreasing fluorescent signal was then measured through Ex. 480 ± 10 nm and Em. 525 ± 10 nm (Gain 60). The time point at which the fluorescent signal reached the basal level was then used to assess the stability of the nanopore. To quantify EDTA quench rates, the developing fluorescent signal was fit to **Eq. S2** through non-linear regression.

**Eq. S2** $$\boldsymbol{Y_{(x)} = (Y_0 - Plateau) \times e^{(-K \times x)} + Plateau}$$

The half-life associated with the decaying fluorescent signal $T_{½}$ = ln (2) / K was used to quantify EDTA quench rates.
For microfluidic analysis, the design of the microfluidic incubation chamber for cultivating *E. coli* was adopted from previous studies[9]. Briefly, individual colonies were used to inoculate 2 mL lysogeny broth (LB) supplemented with 100 µg/mL AMP and 25 µg mg/mL CHL and grown overnight in culture tubes at 37 ˚C and 180 rpm. The following day, 200 µL of this overnight suspension was used to inoculate 10 mL LB supplemented with 100 µg/mL AMP and 25 µg mg/mL CHL, and grown for 2 hours at 37 °C and 180 rpm. The cells were then centrifuged at 11,000 × g for 1 min and washed once with fresh LB medium. After a second centrifugation step at 11,000 × g for 1 min cells were resuspended in 100 µL sterile filtered LB supplemented with 100 µg/mL AMP and 25 µg mg/mL CHL. This suspension was used to seed a microfluidic chamber (manufactured by Wunderlichips GmbH, Zürich, Switzerland). Cells were grown at 37 °C and a flow rate of 4 µL/min controlled by a Flow EZ device (Fluigent) until a sufficient cell count was reached. The LB medium supplemented with 100 µg/mL AMP and 25 µg mg/mL CHL was then changed to LB medium supplemented with 100 µg/mL AMP, 25 µg mg/mL CHL and 25 mM sodium propionate pH 8.0 to induce the G-GECO expression at 30 °C and a flow rate of 4 µL/min. After 110 minutes, the medium was changed again to LB medium supplemented with 100 µg/mL AMP, 25 µg mg/mL CHL, 25 mM sodium propionate and 0.5 mM IPTG to induce nanopore expression. The developing fluorescent signal was monitored by means of a wide-field fluorescence microscope with a 480 ± 5 nm ex. laser (Huebner

GmbH) with appropriate filters F58-019 (Dichroic Mirror) and em. F57-019 GFP/mCherry Dualband filter (AHF Analysentechnik).

*Chemical Synthesis of $S^{21}$68 Holin Variants*

The $S^{21}$ holin variants, $S^{21}$68, $S^{21}$10-68, $S^{21}$24-68 und $S^{21}$30-68 peptides were synthesized on an INTAVIS MultiPep CF peptide synthesizer using a standard Fmoc-SPPS protocol on TentaGel® rink amide resin with a loading capacity of 0.21 mmol/g and *N,N,N′,N′*-tetramethyl-*O*-(1*H*-benzotriazol-1-yl)uronium hexafluorophosphate (HBTU, 5 equivalents relative to loading capacity) as a coupling reagent. The base 4-Methylmorpholine (NMM) in DMF was used in 2-fold excess to amino acids and coupling reagent. Each amino acid was attached by coupling once for 40 min. A real-time monitoring of coupling step was performed, and additional coupling was added automatically when the first coupling was incomplete. The Fmoc- group deprotection was carried out twice by 20% piperidine in DMF. All coupling and Fmoc-deprotection steps were followed by intensive washing of the resin with DMF and dichloromethane. For the $S^{21}$68 peptide, pseudoprolines were inserted as described earlier to facilitate the synthesis[10]. At positions 15-16 and 40-41 Gly-Ser(Psi(Me,Me)pro)-OH, at position 50-51 Leu-Thr(Psi(Me,Me)pro)-OH, at position 28-29 Val-Ser(Psi(Me,Me)pro)-OH and at position 4-5 Ile-Ser(Psi(Me,Me)pro)-OH were inserted. Furthermore, co-solvent N-methylpyrrolidone (NMP) (10% v/v) was added to the coupling mixture and capping by 5% acetic anhydride ($Ac_2O$) in DMF was performed after every $10^{th}$ coupling cycle. All peptides were cleaved from the resin as described previously[11].

Peptides were purified by high-performance liquid chromatography (RP-HPLC) employing a Waters 1525 binary pump and a Waters 2998 PDA detector on a customized Waters 600 module equipped with a Waters 996 PDA detector (Waters, Milford, MA, USA). The gradient elution system was 0.1% trifluoracetic acid (TFA) in water (eluent A) and 0.1% TFA in acetonitrile (eluent B). The peptides $S^{21}$10-68, $S^{21}$24-68 und $S^{21}$30-68 were eluted on a MultoKrom® 100-5 C18 column (250 x 20 mm) column with a linear gradient of 25-60% eluent B for 10 min followed by gradient 60-80% eluent B for 60 min with a flow rate of 8 mL/min. Peptide $S^{21}$68 was eluted on a MultoHighBio® 100-5 C4 column (250 x 20 mm) with a linear gradient of 40-90% eluent B in 120 min with a flow rate of 8 mL/min. The peaks were detected at 214 nm. Collected fractions were combined, freeze-dried and stored at -28°C.

The purity of the collected fractions was confirmed by analytical RP-HPLC on a Waters XC e2695 system (Waters, Milford, MA, USA) employing a Waters PDA 2998 diode array detector equipped with an ISERA® C18 (50 x 4.6 mm, 5.0 µm) column. The molecular weights of the purified peptides were confirmed by ESI mass spectrometry on a Waters Synapt G2-Si ESI mass spectrometer equipped with a Waters Acquity UPLC system.

*Electrophysiological Characterisation of $S^{21}68$ Holin Variants*

The pore-forming properties of wildtype $S^{21}68$ and thereof derived holin variants $S^{21}10$-68, $S^{21}24$-68, and $S^{21}30$-68 were functionally characterised in a vertical bilayer set up at room temperature as previously described[12]. Briefly, chambers were connected with Ag/AgCl electrodes to the head-stage of a patch clamp amplifier L/M-EPC-7 (List-Medical). Applied membrane potentials were referenced to the *cis* compartment. Current traces were filtered at 1 kHz and digitized with a sampling frequency of 5 kHz by an A/D-converter LIH 1600 (HEKA Elektronik). Both chambers were filled with 100 mM KCl 10 mM HEPES pH 7.0 and 1,2-diphytanoyl-sn-glycero-3-phosphocholine (DPhPC) bilayers were formed by air bubble technique[12]. Nanopore peptides were dissolved in DMSO with a concentration of 100 mg/mL and stored up to 3 weeks at 4°C. For bilayer measurements 1 µL of the stock solution was diluted in 1000 µL 100 mM KCl 10 mM HEPES pH 7.0 and 5-10 µL were added directly above the bilayer in the *trans* compartment with a Hamilton syringe. Following successful insertion and assembly of nanopore several rounds of a voltage protocol were applied with 100 mV for 5 sec followed by -100 mV for 5 sec.

### Supporting Results

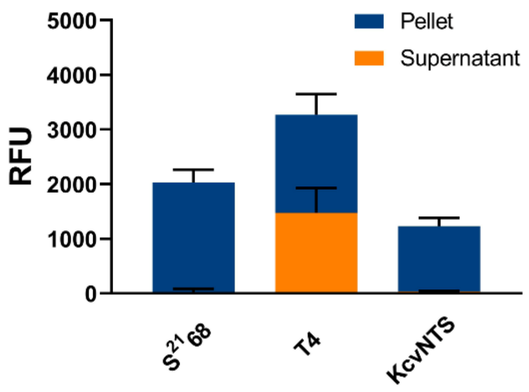


**Fig. S1:** Capacity of different nanopores and ion channels to lyse *E. coli*. Following expression of the $S^{21}68$ holin, the T4 holin or the $K_{CV}$ channel with 0.5 mM IPTG for 3 h in LB medium, cell cultures were spun down and the fluorescent signal measured in the 200 µL supernatant and the pellet (which was resuspended in the same amount of fresh LB medium). For $S^{21}68$ and the $K_{CV}$ channel the fluorescent signal is predominantly located in the pellet indicating that cells remain intact while for the T4 holin close to 50% of the fluorescent signal is in the supernatant indicating that cellular integrity is severely comprised.

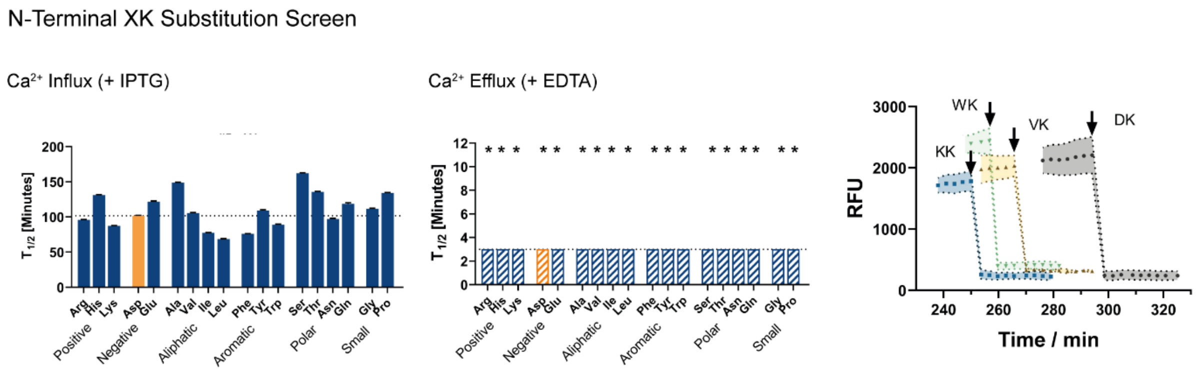


**Fig. S2:** Probing the contribution of N-terminal residues in the assembly and stability of the $S^{21}68$ holin. D2 was substituted with 17 different amino acids but turns out largely insensitive to mutations. The propensity to form stable protein nanopores is assessed in terms of the time $T_{1/2}$ to reach half the maximum signal following addition of IPTG and the half-life $T_{1/2}$ of the fluorescent signal upon addition of EDTA. A set of representative kinetic traces is shown for EDTA quench experiments demonstrating very short decay for D2 substitutions. For mutants denoted with *, the half-life $T_{1/2}$ is less than 3 min and thus below the temporal resolution of the assay. Black arrows denote the time point of adding 5 mM EDTA.

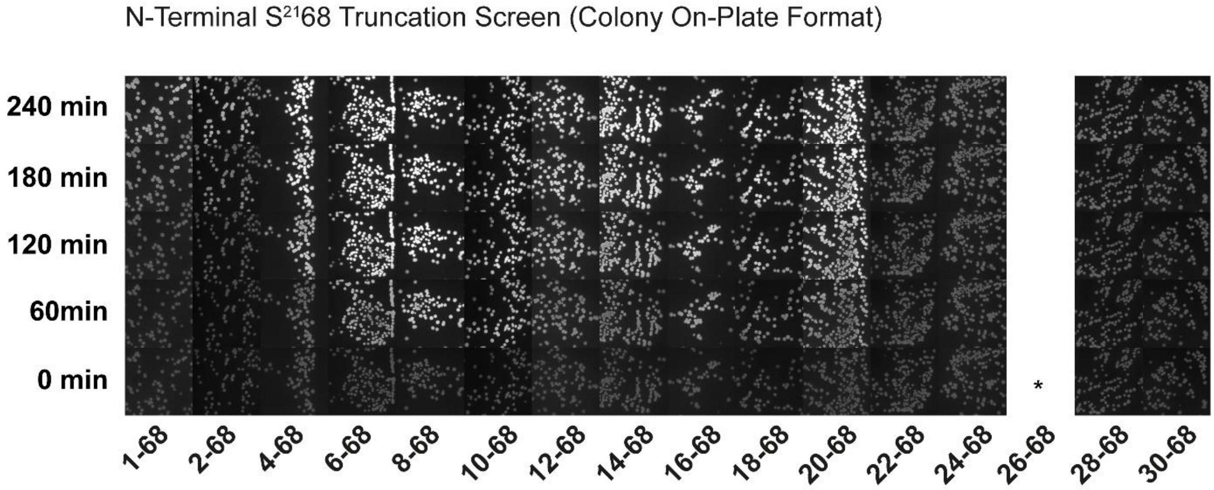


**Fig. S3:** Summary of N-terminal $S^{21}$68 truncations assayed in a colony on-plate format. Crucially, the colony on-plate format recapitulates the quantitative resolution achieved in microtitre plates (see Figure 4B) demonstrating the compatibility of the FuN screen with high-throughput screening formats. The expression of truncation mutants was induced at 0 min with 50 mM IPTG and 500 mM sodium propionate using a spray-painting device and the developing fluorescent signal quantified with a fluorescent imaging reader (E-Box, Vilber).

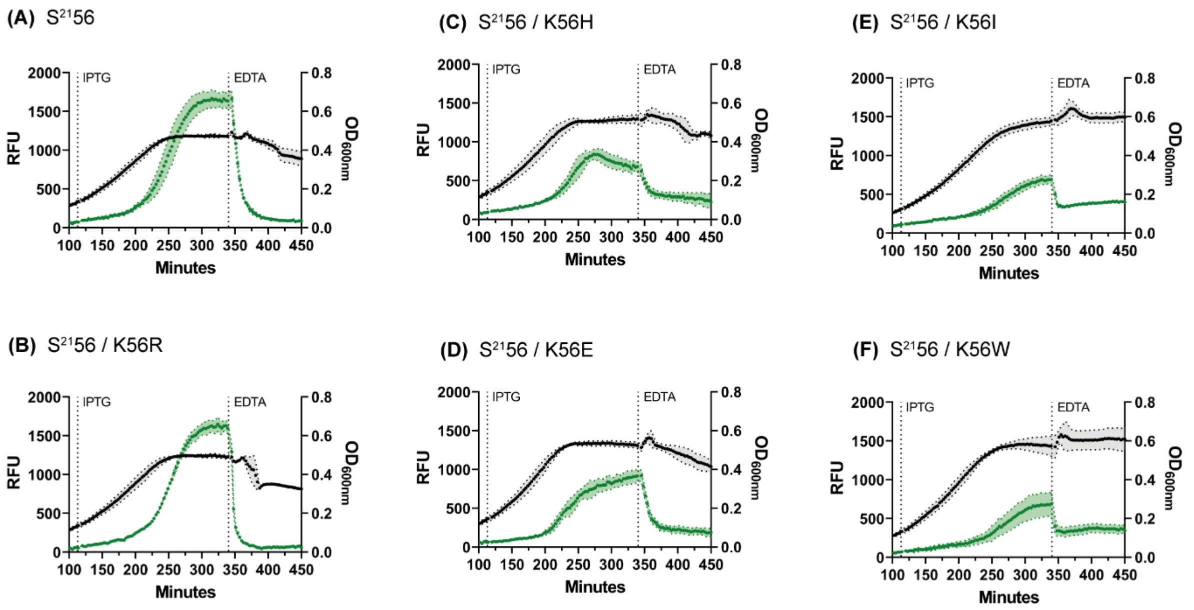


**Fig. S4:** Representative kinetic traces of a select number of $S^{21}56$ truncations with C-terminal K56 substitutions. Only K56R is capable of substituting Lys – even enhancing EDTA quench rates concomitant with its ability to form stronger interactions with phospholipid headgroups – and anchor the C-terminus on the cytoplasmic membrane interface. Alternative substitutions such as titratable His, negatively charged Glu, aliphatic Ile and aromatic Trp show decreasing capacities to form functional nanopores.

### SI References

(1) Gibson, D. G.; Young, L.; Chuang, R.-Y.; Venter, J. C.; Hutchison, C. a; Smith, H. O.; Iii, C. A. H.; America, N. Enzymatic Assembly of DNA Molecules up to Several Hundred Kilobases. *Nat. Methods* **2009**, *6* (5), 343–345.

(2) Gräwe, A.; Ranglack, J.; Weyrich, A.; Stein, V. IFLinkC : An Iterative Functional Linker Cloning Strategy for the Combinatorial Assembly and Recombination of Linker Peptides with Functional Domains. *Nucleic Acids Res.* **2020**, 1–11.

(3) Lee, S. K.; Keasling, J. D. A Propionate-Inducible Expression System for Enteric Bacteria. *Appl. Environ. Microbiol.* **2005**, *71* (11).

(4) Lee, S. K.; Keasling, J. D. Propionate-Regulated High-Yield Protein Production in Escherichia Coli. *Biotechnol. Bioeng.* **2006**, *93* (5).

(5) Ponchon, L.; Catala, M.; Seijo, B.; El Khouri, M.; Dardel, F.; Nonin-Lecomte, S.; Tisné, C. Co-Expression of RNA-Protein Complexes in Escherichia Coli and Applications to RNA Biology. *Nucleic Acids Res.* **2013**, *41* (15).

(6) Imburgio, D.; Rong, M.; Ma, K.; McAllister, W. T. Studies of Promoter Recognition and Start Site Selection by T7 RNA Polymerase Using a Comprehensive Collection of Promoter Variants. *Biochemistry* **2000**, *39* (34).

(7) Gatti-Lafranconi, P.; Dijkman, W. P.; Devenish, S. R. A.; Hollfelder, F. A Single Mutation in the Core Domain of the Lac Repressor Reduces Leakiness. *Microb. Cell Fact.* **2013**, *12* (1).

(8) Kalko, E. K. V; Dukas, R.; Ratcliffe, J. M.; Teeling, E. C.; Haven, N.; Fattu, J. M.; Bates, M. E.; Simmons, J. a; Riquimaroux, H.; Surlykke, A.; et al. An Expanded Palette of Genetically Encoded Ca2+ Indicators. *Science (80-. ).* **2011**, *333*, 1888–1891.

(9) Santos-Moreno, J.; Tasiudi, E.; Stelling, J.; Schaerli, Y. Multistable and Dynamic CRISPRi-Based Synthetic Circuits. *Nat. Commun.* **2020**.

(10) Mutter, M.; Vuilleumier, S. A Chemical Approach to Protein Design—Template-Assembled Synthetic Proteins (TASP). *Angewandte Chemie International Edition in English*. 1989.

(11) Baumruck, A. C.; Tietze, D.; Steinacker, L. K.; Tietze, A. A. Chemical Synthesis of Membrane Proteins: A Model Study on the Influenza Virus B Proton Channel. *Chem. Sci.* **2018**, *9* (8).

(12) Braun, C. J.; Baer, T.; Moroni, A.; Thiel, G. Pseudo Painting/Air Bubble Technique for Planar Lipid Bilayers. *J. Neurosci. Methods* **2014**, *233*.

(13) Dubendorff, J. W.; Studier, F. W. Creation of a T7 Autogene. Cloning and Expression of the Gene for Bacteriophage T7 RNA Polymerase under Control of Its Cognate

Promoter. *J. Mol. Biol.* **1991**, *219* (1).
(14) Elowitz, M. B.; Leibler, S. A Synthetic Oscillatory Network of Transcriptional Regulators. *Nature* **2000**, *403* (6767), 335–338.
